# Costs dictate strategic investment in dominance interactions

**DOI:** 10.1101/2021.06.02.446695

**Authors:** Tobit Dehnen, Danai Papageorgiou, Brendah Nyaguthii, Wismer Cherono, Julia Penndorf, Neeltje J. Boogert, Damien R. Farine

## Abstract

Dominance is important for access to resources. As dominance interactions are costly, individuals should be strategic in who they interact with. One hypothesis is that individuals should direct costly interactions towards those closest in rank, as they have most to gain—in terms of attaining or maintaining dominance—from winning such interactions. Here, we show that male vulturine guineafowl (*Acryllium vulturinum*), a gregarious species with steep dominance hierarchies, strategically express higher-cost aggressive interactions towards males occupying ranks immediately below themselves in their group’s hierarchy. In contrast, lower-cost aggressive interactions are expressed towards group members further down the hierarchy. By directly evaluating differences in the strategic use of higher- and lower-cost aggressive interactions towards competitors, we show that individuals disproportionately use highest-cost interactions—such as chases—towards males found one to three ranks below themselves. Our results support the hypothesis that the costs associated with different interaction types can determine their expression in social groups with steep dominance hierarchies.

## Introduction

In many group-living species individuals have conflicting interests over the use of limited resources [1], resulting in aggressive interactions [2]. Differences in the ability to win such interactions [3] give rise to group-level patterns known as dominance hierarchies. Individuals’ resulting position in dominance hierarchies can have profound consequences for access to important resources such as food [4,5], preferential roosting positions [6] and reproductive opportunities [7], highlighting the importance of rising to the top of the hierarchy—a challenge faced by individuals across the animal kingdom [8]. However, while dominance interactions are ultimately beneficial to some individuals, they also involve costs, including the depletion of energy reserves [9,10], the time spent engaging in dominance interactions [11] and the risk of substantial injury [12] or predation [13]. Accordingly, to maximise the net benefits of investing in dominance interactions, individuals should be strategic in terms of whom they direct their interactions towards, such that they may attain or maintain their position in the hierarchy at minimal cost to themselves [14–17]. Under such a scenario, aggressive interactions should be directed towards competitors closer in the hierarchy, as the directionality of the dominance relationship is likely to be less well-established, and most vulnerable to change, between closely-matched competitors [16,18]. Indeed, theoretical models confirm that directing interactions towards close competitors can stabilise a hierarchy [19]. Beyond deciding who to interact with, individuals might also have the choice to engage in different types of dominance interactions with different group members. More physically-involved or costly interactions are thought to produce more information regarding dominance relationships [20]. Therefore, we can predict that individuals should direct more high-cost interactions towards conspecifics of similar ranks than expected by chance, and this pattern should weaken as interactions become less costly (e.g. lower-cost aggressive or submissive interactions).

Previous empirical studies suggest that, at least in some species or groups, individuals do indeed interact strategically in relation to the relative ranks of group members. For example, Wright *et al.* [14] found a higher frequency of aggression among male mountain gorillas (*Gorilla beringei beringei*) that are close in rank. Similarly, Hobson & DeDeo [21] found that individuals in captive groups of monk parakeets (*Myiopsitta monachus*) express aggression preferentially towards individuals positioned immediately below themselves in the hierarchy, which has since been termed a ‘close-competitor’ strategy [22]. More recently, Hobson *et al.* [22] evaluated many empirical datasets, finding that many social groups and species use a close-competitor strategy. However, there also appears to be extensive within-species variation, with different groups, populations, or species exhibiting ‘bullying’ or a ‘downwards heuristic’ strategies [22]. Such strategies are represented by patterns of aggression expressed preferentially towards either individuals at the bottom of the hierarchy (bullying) or all lower-ranking individuals equally (downward), respectively [22]. Broadening the scope of species studied therefore raises the question of whether a close competitor strategy is universal or not.

A key question to address next is whether individuals are also strategic about what types of interactions they engage in. We can, for example, expect individuals to direct higher-cost interactions (e.g. aggression that involves physical contact, is energetically costly, or can lead to injury) towards individuals closer in rank, whereas lower-cost interactions (e.g. submissive interactions, where one individual simply signals its subordinate status) might be directed more broadly. If this is the case, then variation in strategies across groups or populations might be the product of the frequency and targets of different categories of dominance interactions varying and the subsequent pooling of such interaction categories. For example, if lower-cost aggression is more commonly expressed than higher-cost fights, but merged with the latter, differences in strategies between higher- and lower-cost interactions may be masked. Partitioning out the expression and targets of interactions involving different costs could therefore reveal the nuanced use of cost-driven interaction strategies, and generate insights into underlying decision-making processes.

In this study, we test for strategic use of dominance interactions in vulturine guineafowl (*Acryllium vulturinum*). While living in large and stable groups [23,24] that move together across the landscape [25], group members also regularly form temporary subgroups, exhibit many types of dominance interactions, and maintain a steep dominance hierarchy [26]. We quantify how males of two social groups express higher-cost aggressive interactions (e.g. chases), lower-cost aggressive interactions (e.g. displacements) and submissive interactions towards conspecifics relative to rank difference. We then explicitly test the prediction that males should disproportionately express highest-cost aggressive interactions (relative to lower-cost aggressive interactions) towards individuals nearest in rank.

## Methods

### Empirical data

Our study is focused on two habituated social groups of individually colour-banded vulturine guineafowl living in the savannah and dry-woodland habitat surrounding Mpala Research Centre (0°17⍰N, 37°52⍰E) in Laikipia County, Kenya.

#### Data collection

We collected data between 6^th^ September 2019 and 10^th^ February 2021, recording the actors and recipients of eight distinct types of dyadic dominance interactions (see Table 1) using an all-occurrence sampling method [27]. Observations typically took place while following social groups (on foot or by car) between 0600 and 0930 or 1500 and 1830. We collected data on 76% of the days within the study period. As groups can sometimes be split into spatially distinct subgroups [26], we also recorded group composition information during data collection. We defined different subgroups as cohesive sets of individuals that are spaced ~50 meters or more apart, but also used cues such as direction of movement and cohesiveness to define subgroups. Group one contained 23 and group two contained 29 males. Because only group one resided within a fenced compound—an area surrounded by electric wires to exclude large mammals without impact the movements of smaller species—we had more opportunities to observe interactions from group one than from group two (which was more challenging to observe).

**Table 1.** Vulturine guineafowl (*Acryllium vulturinum*) exhibit a range of aggressive (1-6) and submissive (7-8) dyadic dominance interactions (see also Papageorgiou & Farine [26]). In each description, A represents the actor, and B the recipient, of the interaction.

|  | Interaction category | Interaction type | Description |
| --- | --- | --- | --- |
| 1 | Higher-cost aggressive | SPI | A grabs B by the tail feathers and spins them around |
| 2 | Higher-cost aggressive | TAI | A grabs B by tail or body feathers |
| 3 | Higher-cost aggressive | CHA | A chases B |
| 4 | Lower-cost aggressive | PEC | A pecks B on the head or on body |
| 5 | Lower-cost aggressive | DIS | A displaces B from foraging or resting (e.g. sand-bathing) spot |
| 6 | Lower-cost aggressive | GAP | A gapes at B (similar to chase display but without chasing). B recedes |
| 7 | Submissive | SUV | A falls to the ground in front of B, often accompanied by a ‘crying call’. Usually observed in chicks, but also between adults when on the move |
| 8 | Submissive | SUB | A performs submissive caress with its body under/around the chest of B |

For our study, we only consider interactions that occurred among males. This is partly because the costs and benefits of expressing dominance interactions likely differ when taking place between sexes as opposed to within sex, and may thus follow different strategies. Further, in our study, male vulturine guineafowl tended to engage in dominance interactions most frequently, which is likely because adult males are dominant over all females in the group [26].

#### Interaction dataset structure

Our dominance interactions dataset comprises winner-loser data, which is consistent with most datasets used in studies of dominance interaction strategies [22]. We then rely on the assumption that the winner will be much more likely to be the actor or initiator of the interaction than the loser, thereby allowing us to label winners as the actors and losers as the recipients. This assumption is likely to be correct for our system for several reasons. First, vulturine guineafowl have one of the steepest hierarchies reported to date, with the probability of the dominant winning an interaction being 90% at a rank difference of one, and much greater than 90% at larger rank differences [26]. This makes the outcomes of an interaction likely to be highly predictable for the individual participants, and individuals should generally avoid initiating aggressive interactions they are likely to lose [16]. Second, vulturine guineafowl rarely compete in ‘bouts’ (i.e. discrete events that are won or lost during a fight [17]), with almost all dominance interactions in our empirical dataset being either of short duration or clearly directional (e.g. chases). In species where individuals compete in bouts, back-and-forth aggression occurs before one individual wins, meaning that the winner is not necessarily the individual that *initiated* the interaction. Over years of careful observation, we have only observed potential bouts in very limited circumstances, involving low-ranking males competing over access to females during the peak of the breeding seasons, representing a small fraction of our dataset. In the discussion, we outline further reasons why our results and interpretations are not sensitive to this assumption.

#### Interaction categories

We *a priori* categorised interaction types into higher- and lower-cost categories according with respect to the actor (similar to [28,29]). This categorisation was done by three co-authors (TD, DP and BN) blind to each other’s choices (and all corresponded exactly). Specifically, SPI, TAI and CHA were categorised as higher cost than PEC, DIS and GAP (see Table 1 for definitions). This is because SPI and TAI involve substantial physical contact, in contrast to DIS and GAP that involve no contact. Furthermore, a CHA interaction can last several seconds while a PEC lasts for only a fraction of a second. Additionally, during CHA interactions the actor accelerates and runs at high speed, sometimes covering a substantial distance, which thus represents a considerable energetic investment relative to ‘lower-cost’ interactions involving little to no movement. While the distinction between these interaction categories is very clear in vulturine guineafowl, other studies may opt for a more quantitative approach for grouping interaction types [30], although we note that doing so could potentially require careful consideration of circularity if interaction networks that map differently onto individuals may in fact be the outcome of different strategies. We omit ‘fights’—as defined previously for vulturine guineafowl [26]—because these are infrequent, have no clear directionality, and typically occur between social groups.

Submissive interactions typically function to signal existing dominance relationships—e.g. ‘pant-grunt’ vocalizations in chimpanzees (*Pan troglodytes*) [31,32]—and likely allow individuals to avoid conflict or receiving aggression. Accordingly, the submissive interactions, SUB and SUV, were both deemed to be of low (or no) cost to the actor, and thus grouped into the least-cost ‘submissive’ category.

### Analytical approach for inferring strategic use of dominance interactions

Inferring whether individuals express dominance interactions strategically towards particular group members, and how this varies between interaction types, requires careful consideration of alternative mechanisms that can generate structure or patterns consistent with hypothesised behavioural strategies. For example, even in seemingly closed social groups (i.e. those with stable membership), individuals can vary in their propensity to be in the same part of the group, or subgroup, for example according to rank [33,34] or age and sex [35,36]. Additionally, modelling work has also linked aggression patterns to rank-related spatial positioning [16]. Thus, non-random within-group spatial organisation represents one likely mechanism that could lead to pairwise interactions between some individuals being over-represented in the interaction dataset (simply because they are more often in close spatial proximity), generating patterns resembling strategic use of interactions.

We build on the method proposed by Hobson & DeDeo [21] and Hobson *et al.* [22] to generate estimates of the relationship between the tendency for individuals to express interactions towards conspecifics vs. their differences in dominance rank. We modify this approach in two important ways. First, we deal with potential circularity in the analysis by introducing a randomised data-splitting approach. Splitting the observational data into two parts allows us to independently estimate rank differences and interaction rates for pairs of individuals (step 1 in Figure 1). By repeating the random splitting in a bootstrapping-like process (step 6 in Figure 1)—where observations are randomly re-allocated to each variable in every run—this procedure also provides an estimation of the uncertainty of the relationship. Second, we include a null model—here a permutation test [37]—to quantify how the observed interaction frequency compares to the expected interaction frequency given opportunity to interact alone (steps 3-5 in Figure 1). We refer to this interaction frequency corrected for opportunity as the ‘tendency to interact’.

**Figure 1.**
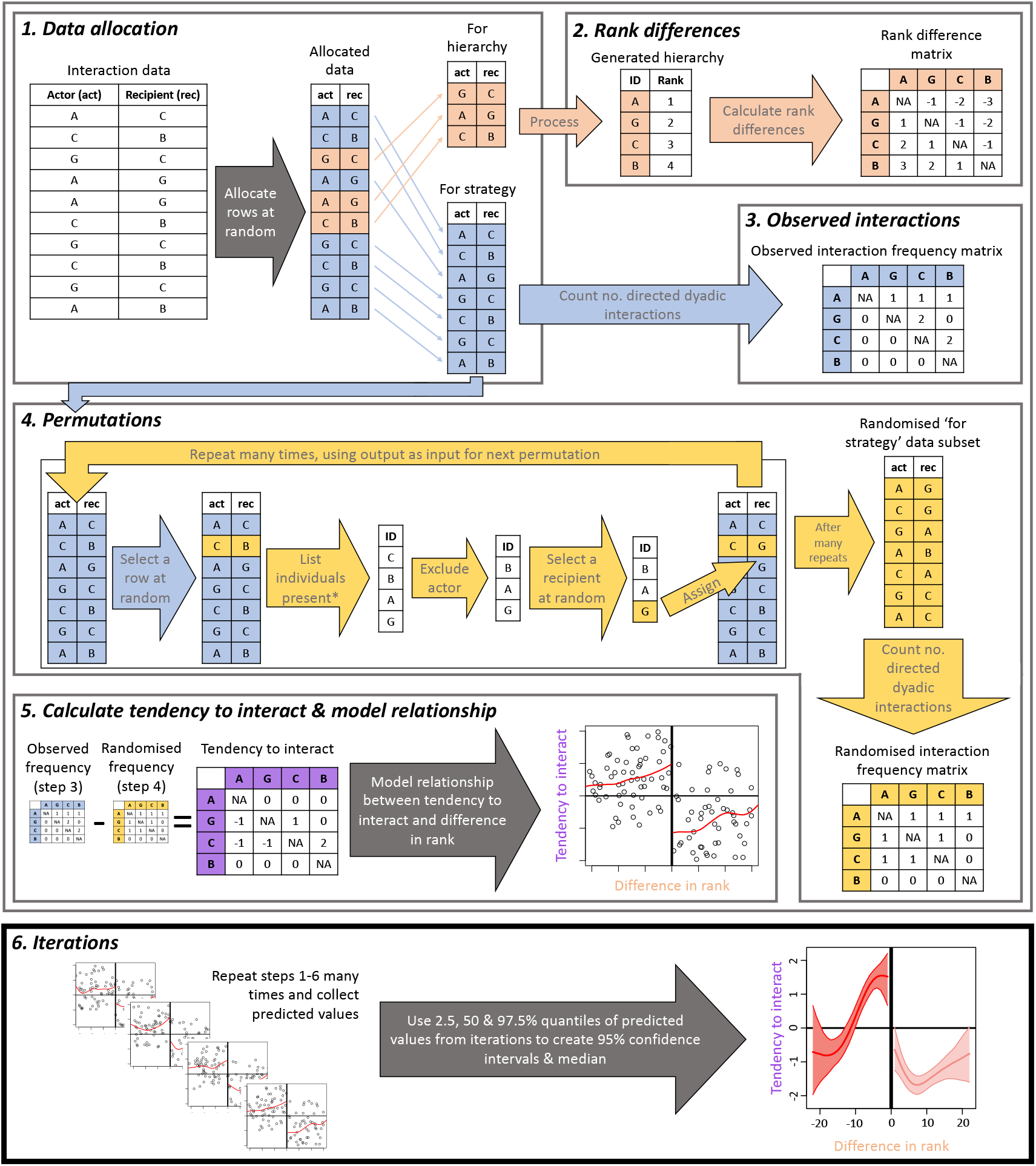
Overview of the analytical steps used in this study. **Step 1:** The dataset of actors (act) and recipients (rec) of interactions (rows) is randomly split into two subsets, with one allocated to generating a hierarchy (orange) and the other to inferring interaction strategies (blue). **Step 2:** A hierarchy is generated and differences in hierarchy position are calculated for each pairwise combination of actors and recipients (dyad). Differences, which can be either in rank or (e.g. Elo) score, are stored in a matrix where the number reflects differences in rank or score. **Step 3:** The sum of directed interactions within each dyad are counted using the ‘for strategy’ (blue) data and stored in a directed matrix. **Step 4:** A permutation procedure—involving the repeated random allocation of interaction recipients from a pool of possible recipients (*based on observation of local group composition at the time the interaction took place)—generates a randomised interaction frequency dataset corresponding to the observed dataset (from step 4). **Step 5:** Differences between the observed and random interaction frequencies are calculated, producing a ‘tendency to interact’ matrix. The relationship between rank/score difference and tendency to interact is then modelled using a method for estimating non-linear relationships (e.g. splines). **Step 6:** Steps 1-5 are repeated many times (e.g. 500), randomly re-allocating different parts of the data to each subset (step 2), recalculating the ‘tendency to interact’ (steps 3 and 4), and storing the predicted values of the model (step 5). The distribution of predicted values are then used to estimate the confidence intervals at each rank difference. The tendency to interact is significantly different to the null expectation, at a given rank difference, when the range between the upper and lower confidence intervals does not overlap 0.

The first step of our approach randomly splits the directed-interactions dataset into two subsets. Given our large observational datasets, and evidence that robust dominance hierarchies can be generated using relatively few observations [38], we use a 30% subset of the data for calculating individuals’ hierarchy positions (and subsequent pairwise positional differences), and the remaining 70% non-overlapping portion of the dataset to infer pairwise interaction rates. Accordingly, the dataset is randomly split to independently estimate the two axes of interest: differences in hierarchy position and directed interaction rates. Future studies, which apply such a data-splitting approach, should make decisions regarding how much data to allocate to each axis by considering the robustness of each inference given the size of the dataset selected.

The second step uses the hierarchy-inference subset of the data to generate a hierarchy. This can be done using a range of methods, and can generate either rank- [39,40] or score-based [41] hierarchies. We use the method proposed by Sánchez-Tójar *et al.* [38], which combines Elo scores with a temporal randomisation procedure to generate a rank-based hierarchy in our main analysis. From these hierarchies, we calculate the difference in rank, whereby the difference in rank is negative if the actor A is higher ranked than the recipient B, and positive otherwise.

The third step is to count the directed interactions across each pairwise combination of actors and recipients (herein a dyad). We choose this option, rather than counting the number of interactions per unit of rank difference, because we naturally expect smaller rank differences to be overrepresented in the data. That is, for N individuals, there are N-1 dyads with a rank difference of 1, N-2 dyads with a rank difference of 2, and so on, with only one dyad having the maximal rank difference of N-1. Thus, simply counting how many interactions occur across all dyads for each unit of rank difference would create a positive linear relationship under the null hypothesis (no strategy). Further, maintaining these data at the dyadic level allows the use of either rank- or score-based estimations of dominance (here we refer to both these as simply ranks). The dyadic interaction count can then be matched up with the difference in rank for that dyad. Note that the number of interactions is generally asymmetric within dyads (individual A may be the actor much more often than individual B).

The fourth step generates an expected pattern of interactions given opportunity. For this, we implement a permutation procedure, which requires data on the local composition of the group (or subgroup) when the interaction took place. Each iteration of the permutation runs as follows:

1. randomly select one interaction and compile a list of all individuals present when the interaction took place,
2. remove the actor (aggressor or submissive individual) from the list, and
3. randomly select an individual from the list as the new recipient of the interaction (this can include the original recipient).

The procedure is run iteratively for a predefined number of permutations (e.g. 100,000), producing a new ‘random’ dataset (i.e. the dataset after the final permutation). Our randomisation procedure differs from previous approaches [21,22] in two ways, based on the specifics of our study. First, we restrict potential recipients to individuals that were present in the subgroup when the interaction took place. Second, we allow for individuals of any rank to be the potential recipient, whereas previous approaches [21,22] restricted potential recipients to those lower in rank. This allows us to describe whether the tendency to interact, at any given rank difference, differed from completely randomly-expressed interactions (see step 5, below), which is essential for comparison across different categories of dominance interactions. By contrast, previous methods [21,22] tested whether the observed strategy differed from a downwards heuristic.

The fifth step generates a measure of tendency to interact that controls for dyadic opportunity to interact. We suggest using the difference between the observed and permuted datasets, i.e. the observed directed interaction count for each dyad minus their expected count from the permuted dataset. Using this measure, zero represents a tendency to interact at the expected frequency, a positive value represents an actor that expresses interactions towards the recipient more often than expected by chance, and negative values represent those that express interactions less often than expected by chance. This measure, estimated for each directed dyad, can be plotted against their difference in hierarchy position, and modelled. We suggest modelling using a method for estimating non-linear relationships, such as splines, because the sum of the tendency-to-interact values is zero, meaning that a linear relationship is only expected when interactions are random or perfectly stratified across ranks.

The above process (steps 1-5) can be repeated many times, each time using different parts of the data allocated to each procedure (step 1), thereby producing an estimate of uncertainty. To estimate the uncertainty, we calculate the 95% range of the predicted tendency to interact-values at set intervals along the difference in hierarchy position-variable from the distribution of splines produced. For differences in hierarchy position in which the 95% range does not overlap with 0, the tendency to interact can be considered as being either significantly greater than expected by chance (when the entire 95% range is above 0) or lower than expected by chance (when the entire 95% range is below 0).

### Testing our approach

We use an agent-based model to confirm that our method is reliable given the parameters of our data, in terms of both group size [23] and hierarchy steepness [26]. In our model, individuals are randomly assigned dominance ranks that remain unchanged throughout each simulation. The model starts by creating *X* subgroups with a given average size *G* drawn from a social group of size *N*. Individuals can be drawn from this social group pool either at random, or can be assorted into subgroups by dominance rank. Thus, we are able to test whether our approach, when inferring tendency to interact, correctly accounts for subgrouping in scenarios where individuals assort non-randomly, here according to rank (as may occur in real animal groups [34]), or when grouped randomly. In the random scenario, each individual has an equal binomial probability 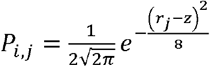 of being drawn in a given subgroup. In the assorted scenario, we first assign a target rank *z* to each subgroup by randomly drawing a number from 1 to *N*, and then setting a binomial probability that each individual *j* with rank *r_j_* is drawn to 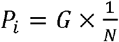. This equation corresponds to the estimated probability density at *r_j_* in a normal distribution with a mean of *z* and a standard deviation of 2. We finally normalise the *P_i,j_* values such that they sum to *G*. For each subgroup, we then set subgroup membership for each individual by drawing a 1 (present) or 0 (absent) using the binomial probabilities *P_i_* or *P_i,j_*, but ensuring that each subgroup has at least two individuals.

For each subgroup, we simulate a single, dyadic interaction by first drawing from the present individuals at random. We then calculate the probability that the more dominant individual would win as *P_win_* = 1 – (1 – *P_w_*)^*r_d_*^, where *P_w_* is the probability that a dominant individual wins given a rank difference of 1, and *r_d_* is the absolute difference in rank (i.e. the difference in rank as a positive number). We then produce an outcome from a binomial draw with the probability of the dominant winning being given by *P_w_* and the probability of the subordinate winning by (1 – *P_w_*), assigning the winner as the actor and the loser as the recipient, matching our observed data.

We evaluate the outputs of both scenarios (random and assorted) using both raw interaction counts and the tendency-to-interact metric. We create *N* = 20 individuals interacting in *X* = 100 subgroups, split the data 70% to calculate strategies, and run 1000 permutations for each iteration of the analysis. We set *P_w_* = 0.9 and *G* = 5.

### Analysis of empirical data

Calculating rank differences: We calculate ranks using the *aniDom* R package v 0.1.4 [38]. We then define rank differences as the rank of the actor minus that of the recipient. For example, if individuals A and B are ranked at 4 and 6, respectively, then the tendency to interact for A to B would be plotted at rank difference=-2 and the tendency to interact for B to A at rank difference=2. We also repeat our main analysis using a score-based hierarchy, by skipping the conversion of Elo scores into ranks, but normalising scores to range from 0 to 1 (as the range of raw Elo scores can differ across different runs of the simulation).

### Robustness of our analyses to methodological decisions

We first test whether hierarchies constructed using each of the three different interaction categories are equivalent. We do this by calculating the correlation between the ranks inferred from each set of interaction categories, for each social group, using the Spearman’s rank correlation coefficient (hereafter *r_s_*). If all three hierarchies are correlated, the interactions in different categories can be assumed to represent the same axis of dominance, confirming that findings across different interaction categories are comparable.

We then test the within-hierarchy robustness of the data by running a repeatability analysis on subsets of the data (i.e. after splitting in step 1 in Figure 1). We randomly select 30% of interactions of a given subset, matching the proportion of interactions used for rank-inference in the main analysis, and use the function *estimate_uncertainty_by_repeatability* from the *aniDom* package [38] to estimate the uncertainty in the hierarchy produced using this 30% subset, producing an *r_s_* value. We repeat this for each interaction category in each group.

#### Testing for strategic use of interactions in the empirical data

We run 500 full iterations of our framework (i.e. 500 repeats of steps 1-5 in Figure 1) for each group and interaction category. In each iteration, the permutation procedure (step 4 in Figure 1) is repeated 100,000 times. We also fit separate splines (step 5) for rank differences above and below zero because interaction strategies towards individuals positioned at relative rank differences of −1 and 1 may differ largely, and forcing a single spline to pass through zero could therefore under-estimate strategies towards close competitors. We use the R function *smooth.spline,* with parameters: df = 3 and lambda = 0.04, to fit all smoothing splines. All analyses are conducted in the statistical environment R v 3.6.2 [42].

Our analysis investigates the strategy within each interaction category, which does not allow us to directly test whether individuals preferentially use one type of interaction versus another according to rank differences. For example, both higher- and lower-cost aggressive interactions might show the same general strategy but to differing extents. We therefore employ Bayes’ rule to calculate the probability of expressing a higher-cost aggressive—as opposed to a lower-cost aggressive—interaction given the difference in rank. The probability of higher-cost aggression *A_h_* in an interaction given a rank difference of *R*, is calculated as 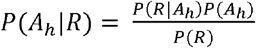. This allows us to evaluate whether there are deviations from the random expression of higher-versus lower-cost aggressive interactions across rank differences. We employ a bootstrapping procedure to estimate the 95% confidence intervals of *P*(*A_h_|R*).

## Results

### Testing our approach

Our agent-based model confirms that our measure of tendency to interact produces expected outcomes both in the presence and absence of strategies (Figure 2), given the structure of our data. By contrast, this is not the case when counting the directed interactions among individuals in groups. These results highlighting the importance of the permutation test to control for potential structure in the data that are caused by processes that are independent of the hypothesis of interest.

**Figure 2.**
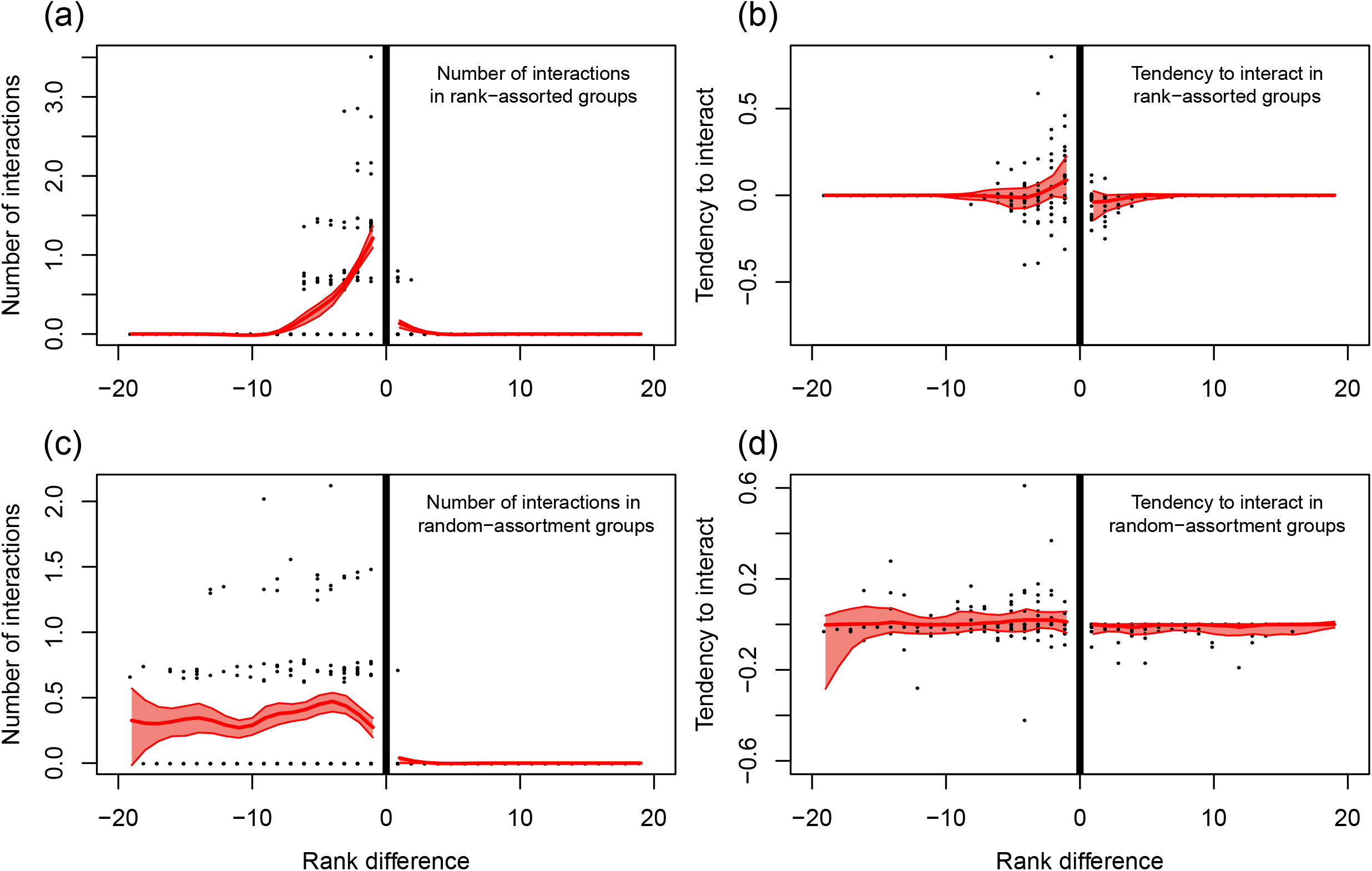
Results from an agent-based model demonstrate the importance of accounting for opportunity to interact in groups reflecting those observed in vulturine guineafowl (*Acryllium vulturinum*). In simulations where individuals are spatially clustered by rank, but interact without any strategy (i.e. at random), this can give the appearance of a close competitor strategy **(a),** potentially leading to spurious inference. Controlling for opportunity to interact using a permutation test can correct for this effect **(b),** with the 95% range of the analyses (red polygon) overlapping 0 across all values of rank difference. When no rank assortment exists, counts of the number of interactions are high across all negative values of rank difference **(c),** making interpretation difficult as the underlying expectation is not explicitly made clear. Accounting for opportunities to interact can confirm that the expression of dominance interactions does not differ from random **(d).**

### Robustness of vulturine guineafowl hierarchies

In total, we recorded 7358 male-male dominance interactions (Table 2). Of these, 25% were classified as ‘higher-cost aggressive’, 28% as ‘lower-cost aggressive’ and 46% as ‘submissive’. The inferred hierarchies are highly robust (all *r_s_* > 0.90, Table 2) when estimated using only 30% of the data for each interaction type within each group. Hierarchies generated using different interaction categories are also highly correlated in both groups (all *r_s_* > 0.95, Figure S1).

**Table 2.** Data summary and within-category hierarchy repeatability for each study group. The *r_s_* value is a Spearman’s rank correlation coefficient, estimating within-dataset hierarchy repeatability as calculated using the function *estimate_uncertainty_by_repeatability* from the *aniDom* package [38].

| Group | Interaction category | no. interactions | Group size | Mean interactions per individual | $r_s$ value |
| --- | --- | --- | --- | --- | --- |
| 1 | Higher-cost aggressive | 1229 | 23 | 53 | 0.96 |
| 1 | Lower-cost aggressive | 1558 | 23 | 68 | 0.93 |
| 1 | Submissive | 2628 | 23 | 114 | 0.95 |
| 2 | Higher-cost aggressive | 627 | 29 | 22 | 0.91 |
| 2 | Lower-cost aggressive | 529 | 29 | 18 | 0.91 |
| 2 | Submissive | 787 | 29 | 27 | 0.93 |

### Interaction strategies

Individuals express aggressive interactions towards individuals below themselves in the hierarchy, but submissive interactions towards individuals above themselves in the hierarchy (Figure 3). This general pattern is expected, as it demonstrates the *existence* of a dominance hierarchy. Individuals in both social groups express significantly more higher-cost aggressive interactions than expected by chance towards group members occupying ranks immediately below themselves, specifically towards individuals positioned 1-10 (peak at 4) and 1-13 (peak at 5) ranks below in groups one and two, respectively. The tendency to express higher-cost interactions with individuals far away in the hierarchy is equal to or less than expected under a random-interactions scenario (Figure 3a,d). In contrast to higher-cost aggressive interactions, individuals in both groups do not express lower-cost aggressive interactions towards their closest competitors more than expected by chance (Figure 3b,e); instead, lower-cost aggressive interactions are preferentially expressed towards individuals slightly further down the hierarchy, specifically towards individuals 2-21 (peak at 7) and 2-14 (peak at 7) ranks below in groups one and two, respectively. At large rank differences, the two groups appear to differ in the strategy inferred for lower-cost aggressive interactions, but we note that the number of dyads and, correspondingly, the number of interactions from which we can draw inference, decreases rapidly as rank difference increases (see Figure S2). Submissive interactions are expressed towards group members positioned higher in the hierarchy at a tendency to interact at, or greater than, expected at random (Figure 3c,f). Our results are consistent when the hierarchy is inferred using Elo scores (standardised to range from 0 to 1) instead of ranks (Figure S3).

**Figure 3.**
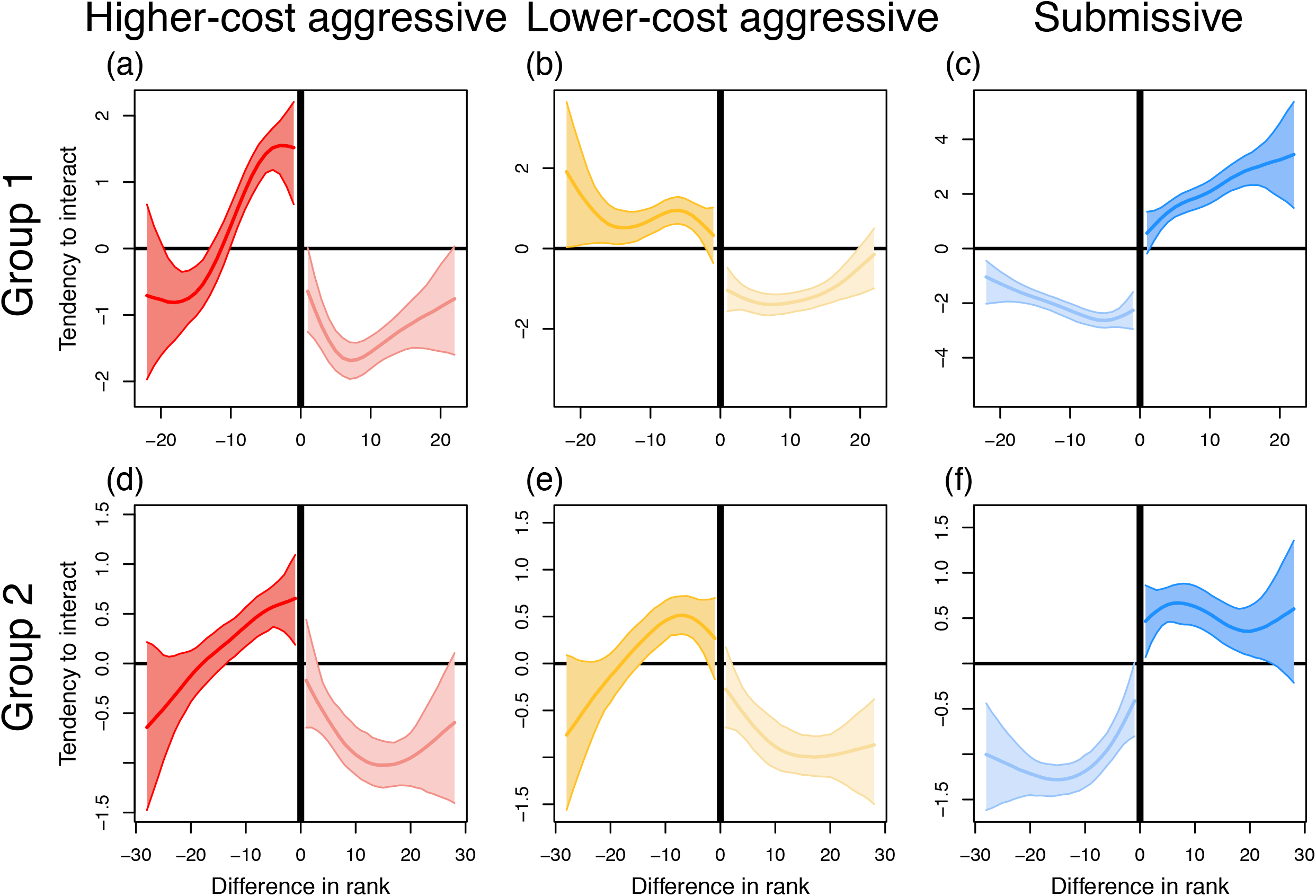
The tendency to interact in relation to relative hierarchy position for three categories of dominance interactions in males of two vulturine guineafowl social groups. Patterns of tendency to interact inferred separately for higher-cost aggressive **(a,d),** lower-cost aggressive **(b,e)** and submissive **(c,f)** interaction categories for social groups one **(a-c)** and two **(e-f).** Each graph shows the median (thick line) tendency to interact and the 95% range (shaded area) of the estimated tendencies (from the repeated data splitting approach described in Fig. 1) plotted against rank difference. A negative difference in rank signifies interactions aimed at lower-ranking individuals, and *vice versa* fora positive difference in rank. The darker side of each graph relates to aggressive and submissive interactions expressed towards lower- and higher-ranking individuals, respectively, and *vice versa* for the lighter side of the graph. Note that the absolute values of tendency to interact depend on the number of observed interactions and are thus not comparable across graphs.

### Aggression decisions

While the above analysis suggests that individuals use cost-based strategies, it does not allow us to identify the relative allocation of different types of interactions according to cost across rank differences. Using Bayes’ rule, we find that there appears to be a tendency for individuals to disproportionately express higher-cost interactions, compared to lower-cost interactions, at very small differences in dominance ranks. At least in group one (Figure 4a), which we had most access to and thereby collected most observations from, individuals express higher-cost aggressive interactions towards group members one to three ranks below themselves significantly more than expected by chance. Group two, which has fewer data (Table 2), has no such pattern, but also much greater uncertainty in the estimates (Figure 4b). Further, the dominant male in group one expresses a surprisingly high number of higher-cost interactions towards the most subordinate individual in the group (i.e. at the largest rank difference).

**Figure 4.**
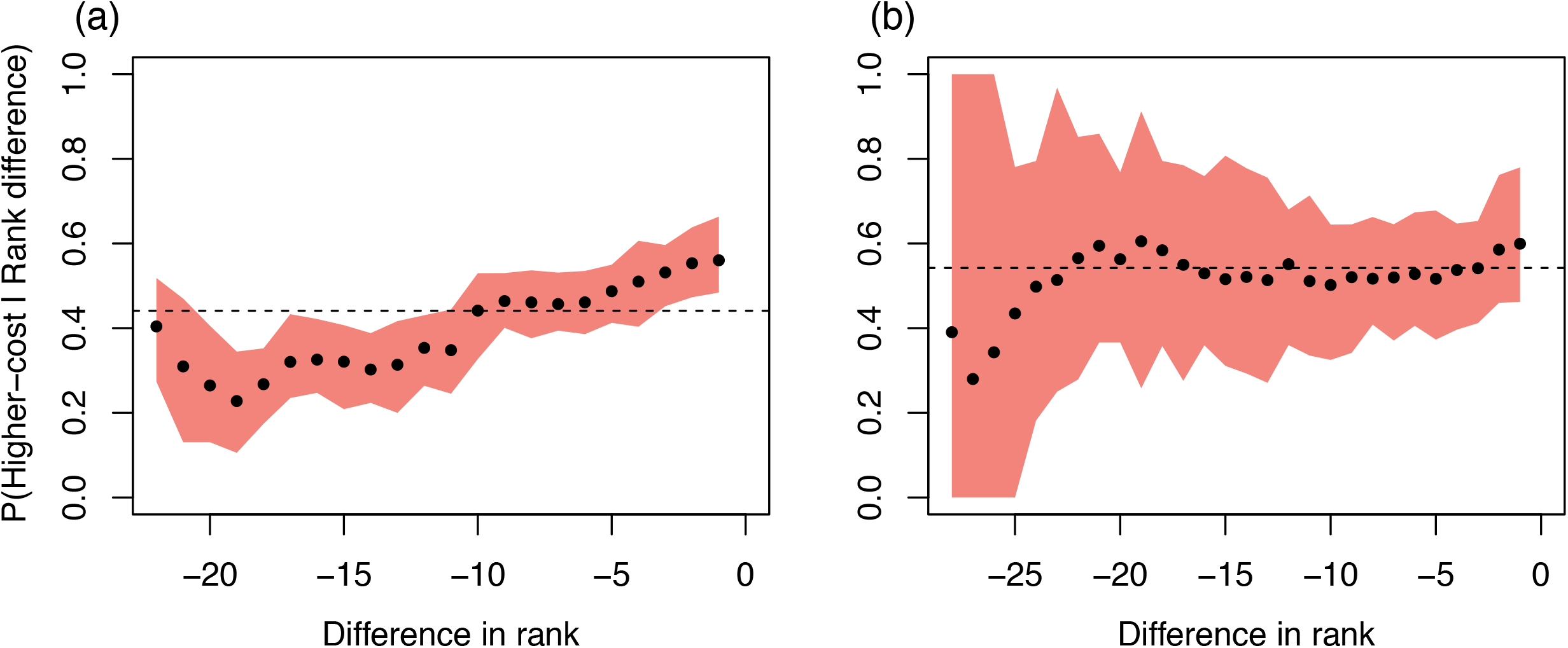
The probability of expressing higher-versus lower-cost aggressive interactions given rank differences in social groups one (a) and two (b). Black circles represent the observed probability of an individual exhibiting higher-cost aggression (versus lower-cost aggression) at a given difference in rank. Shaded areas show the 95% confidence intervals calculated using bootstrapping. The dotted line shows the baseline probability of expressing higher-versus lower-cost aggressive interactions in each group (i.e. *P*(*A_h_*)).

## Discussion

Recent studies have suggested that rank differences may underlie patterns of aggressive interactions in group-living species [21,22]. For example, monk parakeets express aggression towards individuals positioned immediately below themselves at greater frequencies than other group members [21]. Similarly, the frequency of aggressive interactions in both male mountain gorillas [14] and groups of feral dogs (*Canis familiaris*) [43] is higher among individuals close in rank. Here, we extend previous findings by revealing the strategic expression of higher-cost aggressive interactions towards group members positioned close in the hierarchy. This finding is consistent with closely-ranked individuals representing a greater threat of dominance reversals than group members further down the hierarchy, higher-cost interactions potentially providing more information about relative rankings of individuals [20], and evidence that a close-competitor strategy can stabilise dominance hierarchies [19]. Together, our results support the hypothesis that there are multiple axes through which individuals can be strategic in their investments in the context of dominance interactions: the choice of recipient, and the type of interaction.

There are presently two potential limitations to our approach. First, while we account for the dyadic opportunity to interact, patterns suggesting strategic use of higher-cost interactions may be caused by contest escalation among close competitors [17], In vulturine guineafowl, we do not see typical contest escalations, with even higher-cost interactions lacking back-and-forth aggression (i.e. ‘bouts’) and typically being relatively short-lived. Given that contest escalation may involve very subtle behaviours, fine-scale analysis of video sample data could provide insights into the role of escalation in the dominance interactions of vulturine guineafowl. Second, we assume that the winners of aggressive interactions (or losers of submissive interactions) initiate interactions. Due to the steep hierarchies [26] and non-bout aggressive interactions found in vulturine guineafowl, this assumption is likely to be reasonable (see methods for details). While winners are likely initiators in species with steep hierarchies, distinguishing ‘initiators’ from ‘winners’ will likely be more important in systems with shallow hierarchies or in open populations where individuals may be unfamiliar, as a mismatch could produce spurious findings.

Interestingly, if we reverse our assumption of winners being initiators—i.e. we assume that losers initiate interactions—then we would find that individuals express aggressive interactions towards individuals just above them in the hierarchy, with a preference for higher-cost, rather than lower-cost, interactions for the closest competitors. Therefore, we would *still* find that higher-cost interactions are expressed more strategically towards close competitors, but higher-ranking ones. While the biological interpretation may differ, i.e. aggressing those immediately below your rank vs aggressing those immediately above your rank, this still reveals cost-based interaction strategies.

Our findings rely on the assumption that some interactions are more costly than others. In this case, we expect that costlier interactions should be rarer, making it surprising that higher- and lower-cost aggressive interactions are present at similar rates in our dataset. One potential explanation is that higher-cost interactions are more conspicuous and therefore more detectable to a human observer using an all-occurrence sampling method [27,44]. In our dataset, higher-cost aggressive interactions are dominated by chases (CHA interaction code), which happen over distances of metres (and several seconds), making them more obvious to a researcher than the more subtle gapes, displacements or pecks (i.e. GAP, DIS and PEC, the lower-cost aggressive interactions). Our assumed sampling bias is supported by submissive interactions—which are both salient to the observer and should occur most often—being most prevalent in the dataset. An alternative explanation for the high prevalence of interactions in the higher-cost aggression category is that the costs traditionally associated with dominance interactions that involve substantial physical contact or activity, such as energy expenditure [9,10], time investment [11] and the risks of injury [12] or predation [13], may not be so considerable in vulturine guineafowl. While quantifying the costs associated with interaction types warrants further research, our methods are not sensitive to the sampling biases discussed above because they always account for their relative frequency in the data (using permutation tests and through the use of Bayes’ rule).

One interesting, and alternative, perspective concerning the role of costs in predicting investments in dominance interactions is to consider if they are contingent on the dyadic interaction history. For example, theory [45] and recent empirical evidence from vampire bats (*Desmodus rotundus*) [46] suggest that cooperative relationships can emerge by individuals using a raising-the-stakes strategy. Here, individuals may evaluate interaction partners by ‘testing the waters’—using less costly behaviours (such as grooming) that, if reciprocated, would lead to investment in higher-cost behaviours (such as food sharing). The conceptual foundations of such a model are likely to be informative for the study of other types of interactions, such as pair-bond formation [47] and dominance relationships. In the case of the latter, consider two individuals that are relatively unfamiliar, so neither individual holds recent information regarding their relative competitive abilities. One individual might first ‘test the waters’ by approaching, or encroach on the space of, the other. If the outcome of this interaction is unclear, then the next time they interact the individuals) might invest in a higher-cost interaction. Such patterns would be consistent with mutual assessment models [20], such as the sequential assessment model [48], although existing models focus on escalations within a given dominance interaction as opposed to repeated interactions that could take place over weeks or months. Once information about relative competitive ability is available to both individuals, then future interactions may simply omit the escalation process, and jump immediately to the interaction type (usually those that are more costly) that is necessary to keep the information about relative hierarchy position clear. Such a process would generate patterns such as those that we observe in vulturine guineafowl. How dominance relationships become established, or the mechanisms underlying how individuals acquire the information that they then use when expressing interactions strategically, is a promising area for future research.

Our findings add to the growing evidence for group-level dominance interaction strategies across diverse species [14,21,22], and extend the current understanding of such strategies by demonstrating that individuals may use interactions strategically according to their cost. Yet, there are many further axes of strategies to explore. For example, there may be seasonal variation in the importance of dominance rank—such as when food is scarce or when competition for mates is high—that could modulate strategies. Further, studies of dominance interaction strategies, including ours, thus far consider only group-level patterns. Strategies could also vary across individuals, across classes of individuals, or according to individual states, which could be explored using our approach and fitting models informed by other predictors, such as to test the relationship between the dyadic tendency to interact and the sex of the actor. How strategies emerge, the decision rules that underlie their expression, and how these are shaped by features of a given animal society (e.g. the hierarchy steepness) are all open questions ripe for study.

## Acknowledgements

We thank the Mpala Research Centre for logistical support, Kenya Wildlife Service for authorisation to undertake the research (permit KWS-0016-01-21), the National Commission for Science, Technology and Innovation of Kenya (NACOSTI/P/16/3706/6465) and National Environment Management Authority (NEMA/AGR/68/2017) for permits to Kenyan resources and the National Museums of Kenya, and especially Dr. Njoroge, for their ongoing support of our project. In addition, we thank John Ewoi, Mary Waithira and Janet Wangare Kariuki for assistance in the field as well as Lucy Aplin and the Farine lab for discussions about the study.

## Author contributions

DRF and JP conceived the study. TD, DP, JP, NJB and DRF designed the study. BN, WC and DP collected the data. DRF wrote the initial code and TD analysed the data. TD, NJB and DRF drafted the manuscript and all authors contributed to the final manuscript.

## Ethics

Field methods were ethically reviewed by the Max Planck Society Ethikrat Committee (2016_13/1).

## Data Accessibility

All data and code to replicate our analyses are available at https://dx.doi.org/10.17617/3.7z

