## Supplemental Figures for "Costs dictate strategic investment in dominance interactions"

### Supplementary Materials

Tobit Dehnen*, Danai Papageorgiou, Brendah Nyaguthii, Wismer Cherono, Julia Penndorf, Neeltje J. Boogert, Damien R. Farine*

### The PDF includes:

Supplementary text

Supplementary figures S1-S3

### Supplementary Text

Results based on Elo scores: As dominance hierarchies can be represented using both ranks and scores, we investigated whether there were discrepancies in our results based on the metric, i.e. rank vs score, used. Repeating our main analyses (Figure 3 in the main text) using Elo scores instead of ranks yielded qualitatively identical results (Figure S1). This suggests that the hierarchy metric used did not influence our findings.

The role of fine-scale spatial associations in driving interaction patterns: Our methodological developments control for differences in opportunity to interact among dyads that arise as a result of spatial subgroupings. However, it is likely that, even within a cohesive group, individuals are not randomly distributed. For example, dominant white-faced capuchins (*Cebus capucinus*) are more often at the front and centre of moving groups than subordinate group members [1]. Similarly, in vervet monkeys (*Chlorocebus pygerythrus*), dominant individuals are centrally located in stationary groups and at the front when on the move [2]. Further, in groups of coatis (*Nasua nasua*), individuals spend more time in close proximity to group members of the same sex and/or age than expected under a random spatial distribution [3]. Accordingly, given that such differences are typically consistent [4], spatial positioning likely dictates individuals' opportunities to interact with all group members. However, the picture may be more complex, as spatial positioning within (sub)groups is likely mediated by avoidance of aggression from conspecifics [3]. Additionally, whether the probability of aggression being directed to a particular group member increases with proximity or is equal among all individuals present, presumably varies between species and is likely related to the types of resources that are contested. This avenue of enquiry therefore requires further study. Nevertheless, when strategies differ between multiple interaction categories, but these are inferred from a single, simultaneously-collected dataset—as in our study—it seems implausible that proximity alone could drive all of the inferred interaction strategies.

### Supplementary Figures

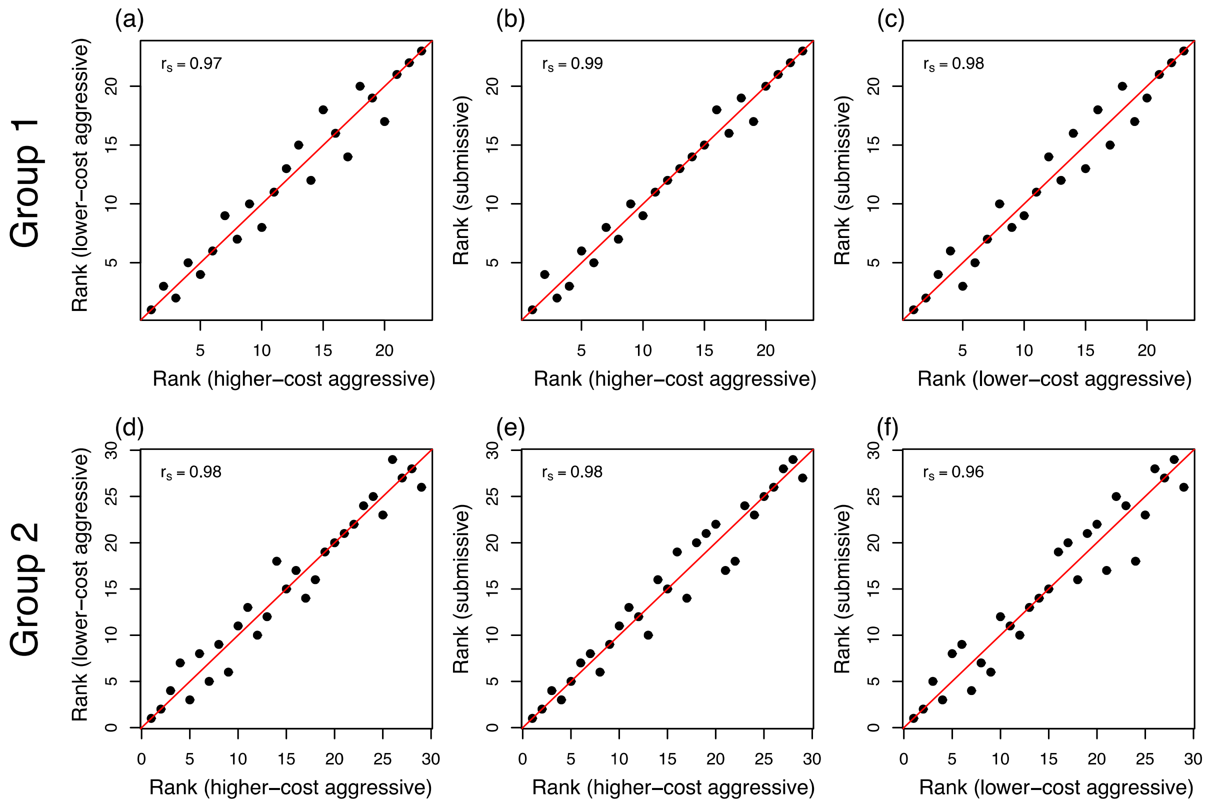


**Figure S1. Correlations between ranks generated using different interaction categories.** The correlation between ranks inferred for all three different combinations of interaction categories using the complete dataset for social groups one **(a-c)** and two **(d-f)**. The Spearman's rank correlation coefficient is printed in the top left corner for each correlation. The y = x line is illustrated in red. The number of interactions in each category used to infer each hierarchy are: higher-cost aggressive: 1229, 627; lower-cost aggressive: 1558, 529; submissive: 2628, 787, respectively, for groups one and two.

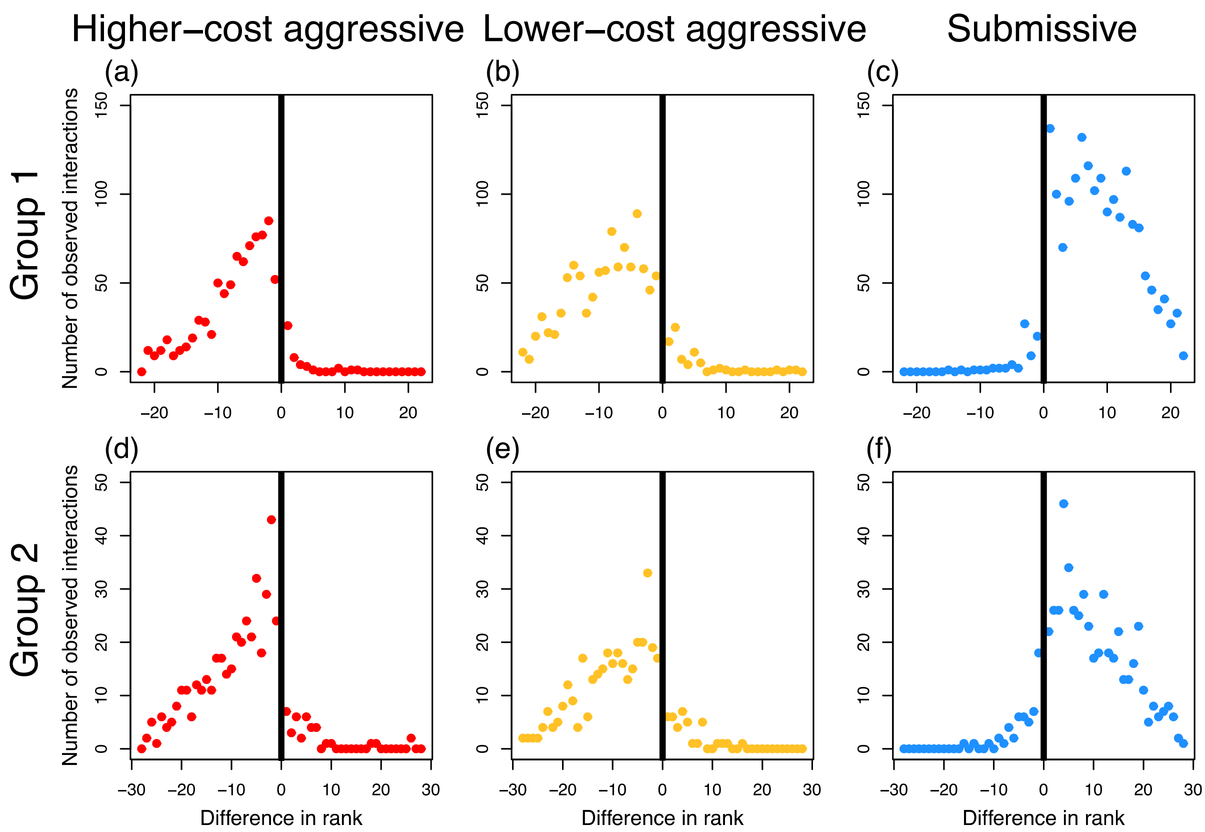


**Figure S2: The total number of interactions at each difference in rank for each 70% data subset.** The number of interactions, summed across all dyads, for each rank difference value for higher-cost aggressive **(a&d),** lower-cost aggressive **(b&e)** and submissive **(c&f)** interactions in social group one **(a-c)** and two **(d-f)**. For each of the six data subsets, the data are split 30-70 (as in the main analysis), for hierarchy inference—i.e. generating ranks and calculating dyadic differences—and counting numbers of interactions, respectively. Therefore, the number of observed interactions in each plot represents a random 70% portion of the data, rather than the entire dataset, for a particular interaction category and group combination. Ranks were generated following the methods of the main analysis.

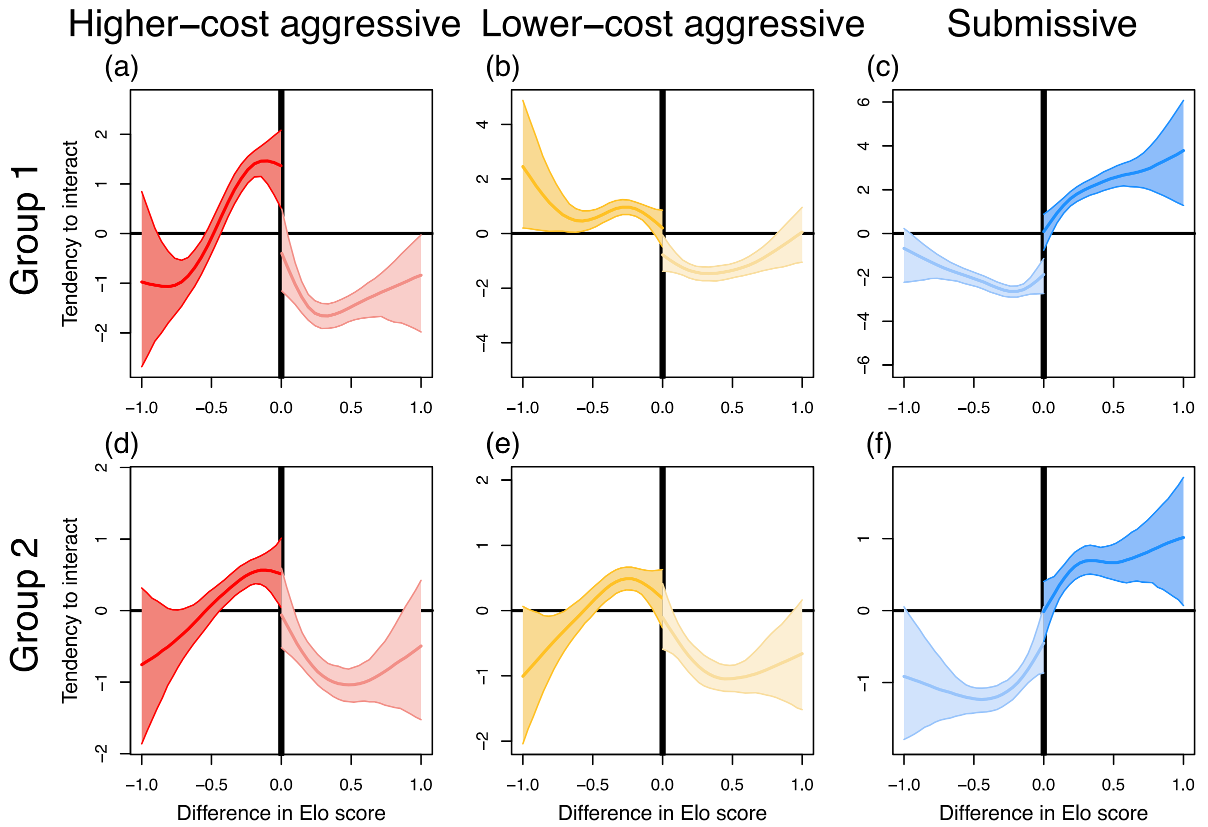


**Figure S3. Repeating the main analysis using ordinal hierarchy.** The pattern of tendency to interact inferred separately for higher-cost aggressive **(a,d)**, lower-cost aggressive **(b,e)** and submissive **(c,f)** interaction categories relative to standardised Elo score for higher-cost aggressive (red), lower-cost aggressive (yellow) and submissive (blue) interaction categories for social groups 1 **(a-c)** and 2 **(e-f)**. Each graph shows the median (thick line) tendency to interact and the 95% range (shaded area) of the estimated tendencies (from the repeated data splitting approach) plotted against difference in Elo score. A negative difference in Elo score signifies interactions aimed towards lower-ranking individuals, and *vice versa* for a positive difference in Elo score. Each side of every graph shows the tendency to interact ranging from the largest (i.e. -1 or 1) to the smallest difference in standardised Elo score among all dyads. Therefore, the median and 95% range may encroach onto the difference in Elo score = 0 line, given that the minimum difference in Elo score may be very small (this is not the case for rank-based hierarchies, where the smallest difference is 1 rank).
